# A Toolbox for Generating Multidimensional 3-D Objects with Fine-Controlled Feature Space: Quaddle 2.0

**DOI:** 10.1101/2024.12.19.629479

**Authors:** Xuan Wen, Leo Malchin, Thilo Womelsdorf

## Abstract

Multidimensional 3D-rendered objects are an important component of vision research and video-gaming applications, but it has remained challenging to parametrically control and efficiently generate those objects. Here, we describe a toolbox for controlling and efficiently generating 3D rendered objects composed of ten separate visual feature dimensions that can be fine-adjusted using python scripts. The toolbox defines objects as multi-dimensional feature vectors with primary dimensions (object body related features), secondary dimensions (head related features) and accessory dimensions (including arms, ears, or beaks). The toolbox interfaces with the freely available Blender software to create objects. The toolbox allows to gradually morph features of multiple feature dimensions, determine the desired feature similarity among objects, and automatize the generation of multiple objects in 3D object and 2D image formats. We document the use of multidimensional objects in a sequence learning task that embeds objects in a 3D-rendered augmented reality environment controlled by the gaming engine unity. Taken together, the toolbox enables the efficient generation of multidimensional objects with fine control of low-level features and higher-level object similarity useful for visual cognitive research and immersive visual environments.

## Introduction

Investigating cognition requires tools that can precisely manipulate and control the features of stimuli presented to participants. Multidimensional, feature-controlled object sets have become indispensable in cognitive research for probing perception, recognition, and categorization processes (Arnott et al., 2008; Biederman & Gerhardstein, 1993; Bowman & Zeithamova, 2020; Chuang et al., 2012; Ghazizadeh et al., 2016; Knutson et al., 2012; Mercer & Duffy, 2015; Pusch et al., 2023; Todd, 2004; Wallraven et al., 2014; Wong et al., 2009). In previous research, Watson et al., 2019 introduced a toolbox that generated 3D-rendered objects with fine user control of up to five different visual feature dimensions (Watson et al., 2019). Multidimensional objects generated with this toolbox, so called Quaddle objects, have been effectively utilized to explore various aspects of cognitive processing, such as object recognition, discrimination, visual search, attentional set shifting and feature learning (Banaie Boroujeni et al., 2022; Boroujeni et al., 2022; Hassani et al., 2023; Kemp et al., 2024; Womelsdorf, Thomas, et al., 2021; Yazdanpanah et al., 2024). Moreover, 3D-rendered multidimensional objects are used in cognitive research using augmented reality (AR) and virtual reality (VR) that provide immersive environments with the aim to mimic real-world experiences (Smith, 2019). For these dynamic and sometimes interactive settings 3D rendering of objects can enhance task performance and have been used in contexts studying object identification, spatial manipulation and navigation, object memory recall and learning (Corriveau Lecavalier et al., 2020; Howett et al., 2019; McIntire et al., 2014).

Despite the usefulness of 3D rendered objects in the described settings, researchers face challenges in generating well controlled 3D objects due to technical constraints and the complexity associated with 3D modeling software. Addressing this situation, we introduce the Quaddle 2.0 toolbox, an evolution of the original Quaddle toolbox designed to enhance utility and flexibility of object generation for dynamic visual and cognitive science studies and video-game contexts (Watson et al., 2019). Quaddle 2.0 expands upon its predecessor by increasing the available feature space (from five to ten visual feature dimensions) (**Figure 1A**), while still allowing generating simpler Quaddle 1.0 objects that has less feature dimensions (**Figure 1B**). In addition, the Quaddle 2.0 toolbox newly establishes the use of Blender -a free, open-source 3D creation suite compatible with macOS and Windows – to interface flexibly with python scripts, allowing an efficient automatized control over low-level object features (gradual feature value changes) and higher-level 3D object characteristics (e.g. light viewing angle).

**Figure 1.**
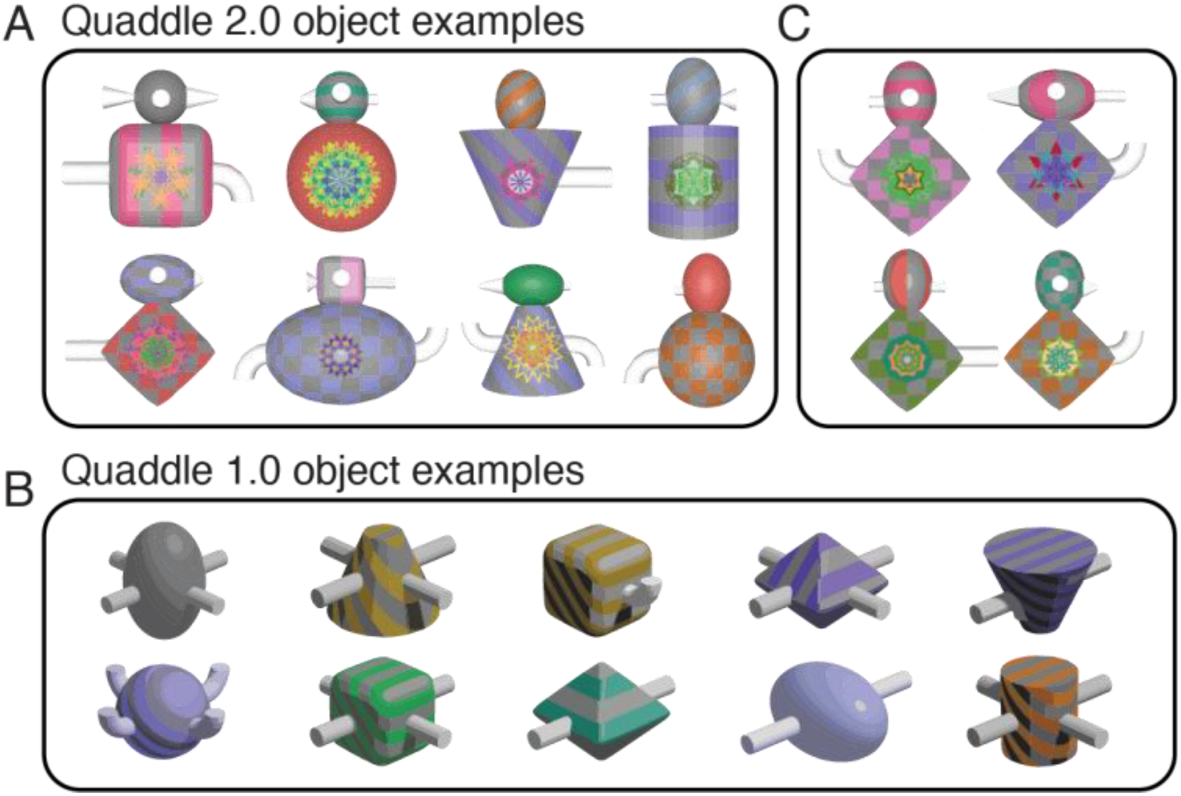
Examples of Quaddle objects. (**A**) Quaddle 2.0 objects varying in ten different feature dimensions. (**B**) Simpler objects that vary in features of four dimensions (body shape, arm angle, body colors, body pattern) shows the toolbox allows reproducing Quaddle 1.0 objects that were generated with a different software rendering suite (Studio DS Max). (**B**) Quaddle 2.0 objects that are more similar to each other by setting the similarity score > 8.

Here, we (i) detail the Quaddle 2.0 framework, (ii) introduce the scripting pipeline for generating large object sets, (iii) illustrate how users can control the feature similarity among objects, (iv) demonstrate the gradual morphing of multiple object features, and (v) showcase the use of Quaddle 2.0 objects in an experimental environment programmed with the unity game engine to evaluate how non-human primates learn the sequential ordering of visual objects. Example code (in python and MATLAB) and a documentation with detailed instructions for using and customizing Quaddle 2.0 objects is freely available online (see **Appendix**).

## Method

### Composition of Quaddle 2.0 objects

Quaddle 2.0 objects vary in features of ten pre-configured object dimensions (**Figure 2**). The default object contains five main parts: head, body, arm, ear, and beak. The head and body can be covered with surface color and patterns, and the body can have pre-configured, colorful fractal images projected on its center. The default arm, ear and beak are colored white and can be adjusted to have different length, shape and bending angle. Arm, ear and beak are considered minor feature dimensions that can be omitted from the objects (**Figure 2**).

**Figure 2.**
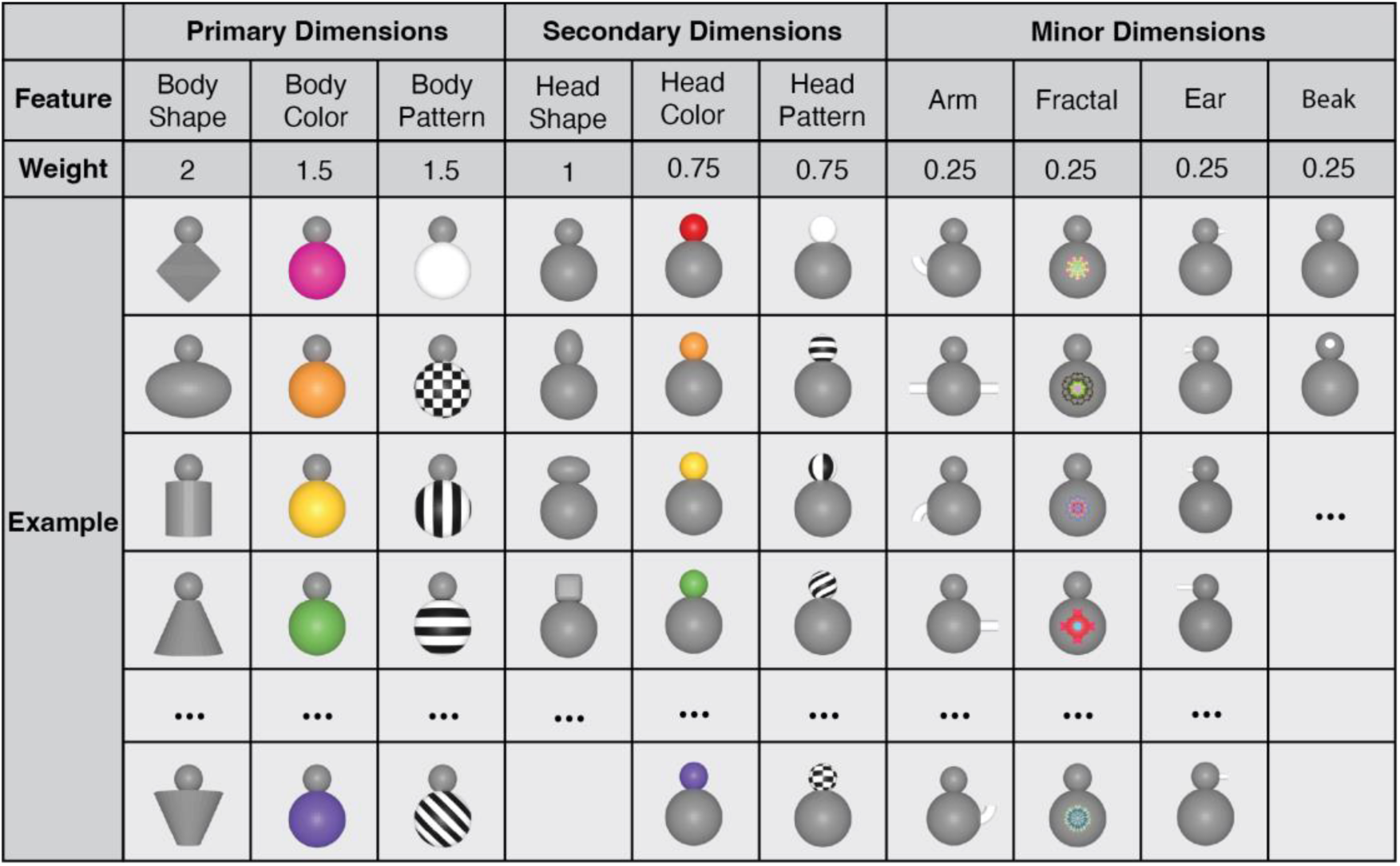
Object dimensions and example feature values pre-configured in the Quaddle 2.0 toolbox. Feature dimensions are categorized into primary dimensions (body-related), secondary dimensions (head-related), and minor dimensions (accessories), organized by size and front-view visibility. The ‘weight’ row denotes how a feature dimension is weighted when calculating the similarity of objects (*see* similarity score calculations).

### Generating 3D objects using lists of pre-configured object features

Quaddle 2.0 objects are rendered using Blender (blender.org), a free and open-source 3D modeling software suite. Objects generated for figures in this article used Blender software version 3.4.0. Generating Quaddle 2.0 objects follows a structured pipeline implemented through a series of Python scripts within a Blender project file as outlined in **Figure 3**. All Python scripts are freely available online (*see* **Appendix**). The process begins with user-specified information about the desired feature dimensions and feature valuer per dimension, which are provided via a command-line tool or in a text file. The input is parsed using a parser function, which extracts the values for each visual feature of the different object dimensions. The core of the generation process involves several scripts that sequentially build the Quaddle as surveyed in **Figure 2**. Each component of the Quaddle is generated through a series of modular functions. The process begins with the creation of the main body, followed by the attachment of the head. Subsequently, additional functions append the ears to the head, add arms to the body, and attach a beak to the head. This modular approach allows for precise control over each component of the Quaddle, enabling systematic manipulation of individual features while maintaining overall structural consistency. The body of the Quaddle object is initially generated from the default mesh shapes in Blender, including sphere, cube, cone, and cylinder, and then molded into the user-specified desired body shape. The arms, beak and ears are initially generated as straight cylinders and molded into the desired angle, shape, and length, and which allows gradual morphing of object features by specifying intermediary values in bended angle, size of shape and length. In the feature set pre-configured in the Quaddle 2.0 toolbox, each Quaddle has at most two arms, two ears and one beak (**Figure 4**). There are three angles for arm (bend up, bend down, straight), three shape for ear (straight, pointy and blunt), and three lengths for ear (regular, long, short). Different sub-features can be combined freely. Additional feature dimensions and feature values can be realized by modifying the pre-configured python code.

**Figure 3.**
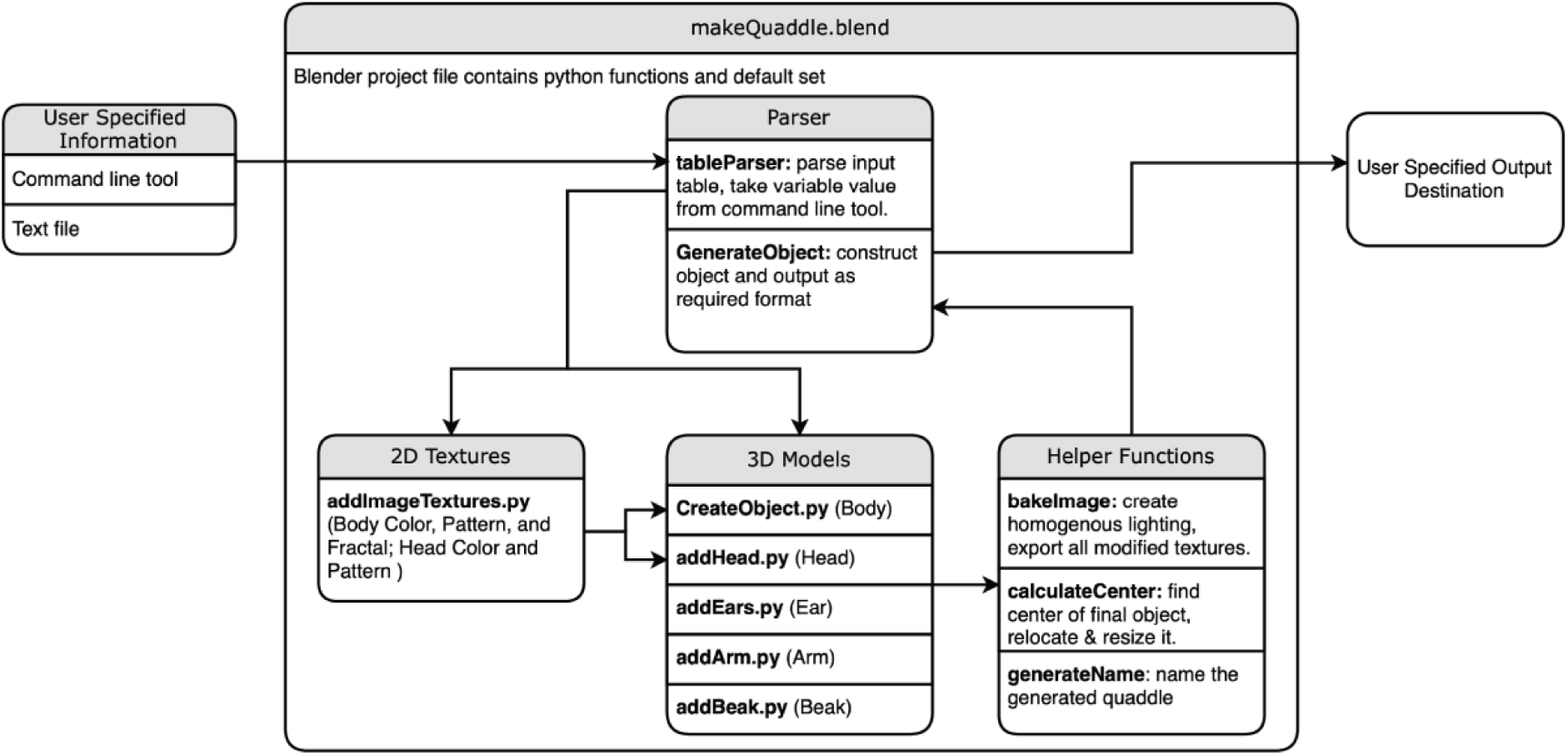
Generation pipeline for Quaddle 2.0 objects. The automated pipeline for generating Quaddle models using the *makeQuaddle.blend* Blender project file, which contains predefined Python functions. User-specified input (either via a command-line tool or a text file) is parsed by *tableParser.py*, which extracts the parameters used to generate the object with the script *GenerateObject.py*. The main pipeline splits into two key branches: (1) **2D Textures**, where functions apply body and head colors, patterns, and body fractals; and (2) **3D Models**, where scripts build the respective morphological elements. **Helper functions** handle lighting, object positioning, and naming before exporting to the user-specified destination.

**Figure 4.**
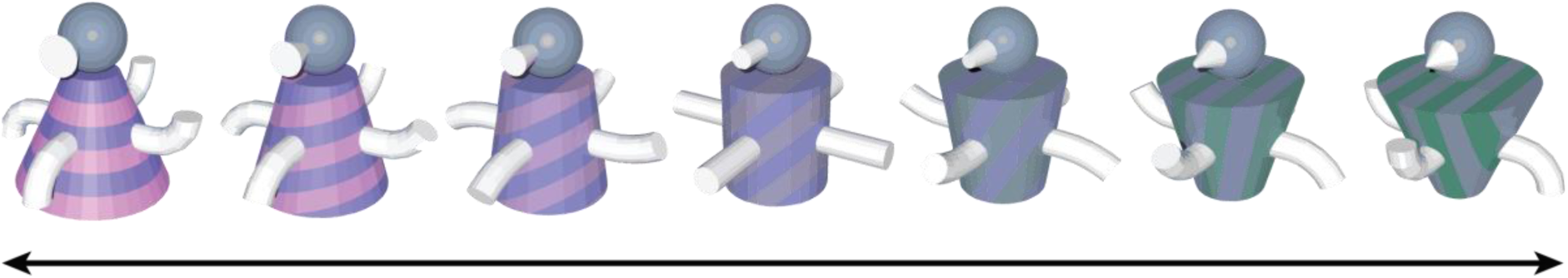
Illustration of gradual morphological changes between feature dimensions using the Quaddle 2.0 toolbox. The objects transition in body shape (from conical to cylindrical to rectangular), arm angles and orientations, color and surface pattern (from pink and purple horizontal stripes to green and blue vertical stripes), and beak shape (blunt to pointy)

### Generating 2-D textures and surface features

Following the structural assembly, 2D textures are applied to the 3D model. This script *addImageTextures.py* manages the application of body color, pattern, and fractal designs, as well as head color and pattern. The surface colors and patterns are imported from PNG files. The default gray color is the same for all objects, whereas the other colors are chosen within the CIE L*c* h* space such that the L* and c* values (luminance and saturation, respectively) are held constant, but h* values (hue) vary by 30°, meaning that there is a small difference in hue between the two colors, but not in the other components of the colors. The texturing process employs a UV project modifier, a specialized tool in Blender that projects 2D images onto the 3D object’s surface. This modifier projects images from the front of the object, minimizing distortion when viewed frontally and ensuring symmetrical pattern application on both front and back. Two separate UV project modifiers are utilized for background patterns and surface fractals separately, each with a distinct projector object for optimal surface coverage. The process involves creating a material with a node-based shader network. Two image textures are incorporated and combined using a mixing operation, allowing for complex surface patterns. The first texture applies directly to the object surface, while the second serves as both a secondary color source and a blending mask. This approach prevents distortions of frontal views, facilitates symmetrical patterns, and allows control over texture placement and scale. By manipulating UV maps and projector properties, the process achieves various texture effects, from uniform coverage to complex, position-dependent patterns. The projection method can be similarly applied across different object geometries, which allows using the same complex visual pattern across a diverse set of object shapes.

The Quaddle 2.0 toolbox allows projecting fractal images onto the object body or to render various body surface patterns. Fractal images are used because they provide a large and abstract image space that is unfamiliar to experimental subject. The fractals were generated following previously published methods (Ghazizadeh et al., 2016) using point-symmetrical polygons, where parameters such as size, edges, and color were randomly chosen to create distinct visual shapes. This method ensures that a large number of distinguishable fractals are available.

With regard to patterns for the body surface of objects, the Quaddle 2.0 toolbox has five pre-configured patterns available: *solid* (default), *horizontal stripes*, *diagonal stripes*, *vertical stripes*, and *grid*. Each pattern is generated as a 1200x1200 PNG file, with stripe widths set to 150 pixels to ensure the patterns and their associated colors remain clear and distinguishable in the final stimuli. While users can select any combination of colors for the patterns from the prepared asset folder, the current code defaults to combining a neutral grey with a user-specified secondary color (for examples, see **Figure 1**). This design ensures that the colors are well visible and have apparent contrast in the patterned image.

### Refinement and exporting 3D rendered objects

Post-processing steps are then executed to refine the generated Quaddle objects. These steps include creating homogeneous lighting conditions and exporting all modified textures, ensuring consistent visual quality across stimuli. The process also involves locating the center of the final object, repositioning and resizing it as necessary to maintain standardized dimensions across all generated Quaddle objects. Additionally, a unique identifier is assigned to each generated object, facilitating organized data management in subsequent experimental procedures.

The final stage constructs the complete object based on all specified parameters and outputting it in the required format to a user-defined destination. The Blender Python script that can read multiple object definitions at once and create large sets of object, exporting them either as image files (e.g., PNG), or 3D shape files (e.g., FBX, GLTF) with any specified distance, height, and rotation. The default output centers the object at the middle points of height and width of each object. The camera distance is adjusted based on the size of the objects with the aim to maintain a consistent apparent size across objects.

### Controlling the similarity of multidimensional objects

Quaddle 2.0 objects can share variable numbers of features, which we exploit by calculating a similarity score between objects. Users can set a similarity score in the *make_quaddle.m* MATLAB script when generating two or more objects. Similarity is calculated by comparing the values of each feature between two objects, with different weights assigned to each dimension. Visually more prominent object dimensions weight strongest (weights of body shape: 2; body color: 1.5, body pattern, 1.5), while less prominent secondary dimensions weight less (head shape, head color and head pattern: weight 0.75), and minor dimensions weight least (Arm, Fractal, Ears, Beak: weight 0.25). Using these feature dimension weights the similarity score consider pairs of Quaddle objects ***Q***_1_ and ***Q***_2_, defined by their dimension ***Q***_1_ = [*q*_11_, *q*_12_, … , *q*_1*n*_] and ***Q***_2_ = [*q*_21_, *q*_22_, … , *q*_2*n*_]. A binary vector indicates whether the values in each dimension are the same is defined as ***B*** = [*b*_1_, *b*_2_, … , *b*_*n*_] where *b*_*i*_ = 1 if *q*_1*i*_ = *q*_2*i*_, else *b*_*i*_ = 0. Weights are predefined for each feature dimension *i* : ***W*** = [*w*_1_, *w*_2_, … , *w*_*n*_] . The Similarity Score S is calculated as*S* = (***W***)^*T*^(***B***) or equivalently as 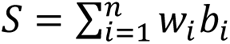
. Using the pre-configured Quaddle 2.0 objects, the similarity score ranges from 0 (most dissimilar) to 11.75 (most similar). We validate the similarity score by comparing the similarity of object pairs as defined by the score with other quantitative similarity estimates (see below and **Figure 5**) and by empirically evaluating the perceptual confusability of objects with high versus low similarity scores (see below and **Figure 6**).

**Figure 5.**
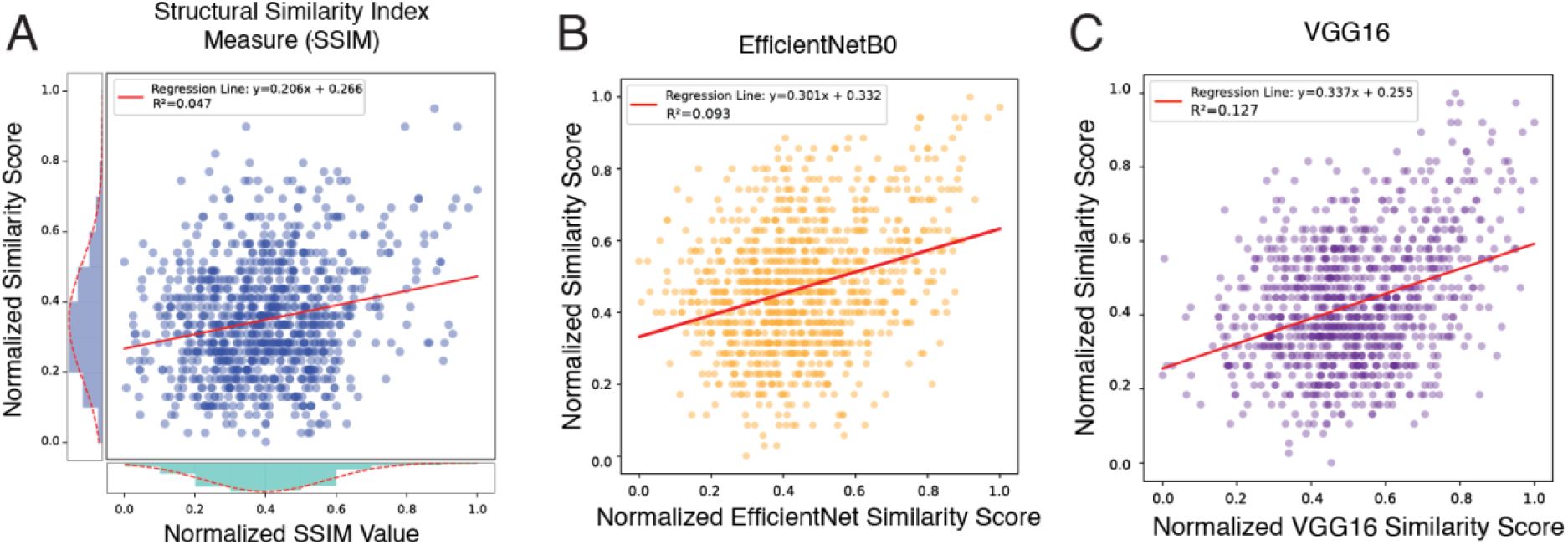
Comparison of object similarity scores. **A:** Scatter plot showing the relationship between normalized SSIM values and similarity scores, including the distribution of both scores. The regression line is y=0.206x+0.266 with *r*^2^=0.047. **B:** Scatter plot illustrating the correlation between normalized EfficientNetB0 similarity values and similarity scores. The regression line is y=0.301x+0.332 with *r*^2^=0.093. **C:** Scatter plot depicting the association between normalized VGG16 similarity values and similarity scores. The regression line is y=0.337x+0.255 with *r*^2^=0.127. Each figure is based on a random draw of 1000 pairs of 300 randomly generated Quaddle images.

**Figure 6.**
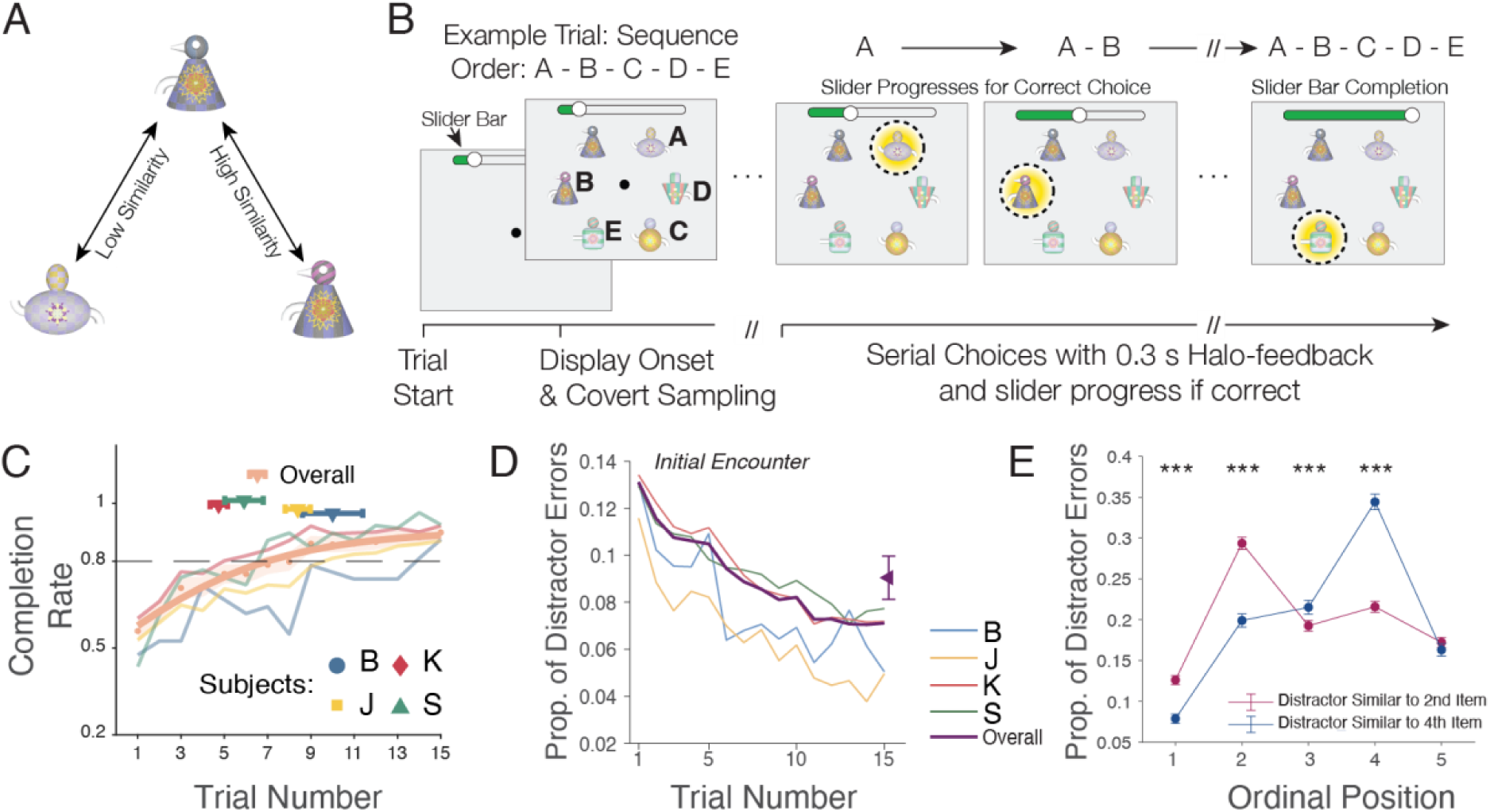
Experimental paradigm and results. A: Illustration of Quaddle 2.0 objects that are perceptually dissimilar (left) and similar (right) as quantified by the similarity score (*see* text for details). **B**: Each trial presented six objects. Monkeys learned to touch five objects in a pre-determined order A-B-C-D-E and avoid a distractor object. A correct choice led to visual feedback (yellow halo) and incremented the slider position on top of the screen. Subjects had 10 maximum number of errors in exploring one sequence. Successfully completing one sequence would lead to fluid rewards. In each trial, distractor object has higher similarity to either 2^ed^ or 4^th^ object in the sequence, compared to lower similarity with other objects. **C:** Learning curves for completing sequences with ≤ 10 errors (y-axis). Symbols show avg. trial and SE to reach 80% completion for each subject and overall. **D:** Proportion of distractor choices across trials. The mean value is shown as the rightmost data point (Mean ± 95% CI: 0.090 ± 0.010). E: Distractor choices at each ordinal position when distractor is similar to object at the 2^nd^ ordinal position and when distractor is similar to object in 4^th^ ordinal position. Two-proportion Z-tests were applied at each ordinal position for comparing the difference. Stars denote sign. level. (Ordinal Position: Before & After Swap, Mean ± 95%CI) 1: 0.13 ± 0.01; 1: 0.08 ± 0.01; 2: 0.29 ± 0.01; 2: 0.20 ± 0.01; 3: 0.19 ± 0.01; 3: 0.22 ± 0.01; 4: 0.22 ± 0.01; 4: 0.34 ± 0.01; 5: 0.17 ± 0.01; 5: 0.16 ± 0.01. Figure panels B and C were adapted from Wen et al., 2024.

### Experimental paradigm for serial ordering of Quaddle objects with controlled feature similarity

To demonstrate a use case for the 3D rendered Quaddle 2.0 objects we generated multiple sets of Quaddle objects and trained four rhesus macaque monkeys to learn pre-defined temporal (serial) orders of five objects in each those object sets in a sequence learning task. We defined the sixth object as a distractor with a high similarity to only one of the other objects to illustrate the distractor is confused specifically with an object that has a high similarity to it as defined by the similarity score. All animal and experimental procedures used in this study complied with the National Institutes of Health Guide for the Care and Use of Laboratory Animals and the Society for Neuroscience Guidelines and Policies and were approved by the Vanderbilt University Institutional Animal Care and Use Committee.

The details of the experimental set-up and the timing of task events are described in detail in (Wen et al., 2024). In brief, four adult male rhesus monkeys performed the sequence learning task in their housing cage using cage-mounted touch screen Kiosk stations (Womelsdorf, Thomas, et al., 2021). Visual display, behavioral response registration, and reward delivery was controlled by the Multi-Task Suite for Experiments (M-USE) (Watson et al., 2023). M-USE is a free software platform with pre-configured tasks (https://m-use.psy.vanderbilt.edu) that displays 3D rendered objects using the Unity3D program (here: Unity version 2020.3.40F1). M-USE displayed the objects on an Elo 2094L 19.5" LCD touchscreen with a refresh rate of 60 Hz and a resolution of 1,920 × 1,080 pixels, rendered at approximately 3 cm on the screen. Experimental sessions tested at least eight unique sequences, each with a unique set of six Quaddle 2.0 objects. Sets of objects were generated by randomly assigning objects different features from up to ten different feature dimensions (see **Figure 1**). Five of the objects were defined to be dissimilar to make them easy to distinguish, while a sixth object was a distractor that was not part of a sequence but shared features with either the object at the second or the fourth serial order (**Figure 6A**). The similarity of the distractor with selected objects of the sequence was used to validate that the similarity of Quaddle 2.0 objects translated into perceptual confusability among objects.

As illustrated in **Figure 6B**, the task presented on each trial six objects equidistant to each other at random locations on a circle equidistant from the center of the screen. Five of the objects were assigned a unique ordinal temporal position in the sequence of objects A-B-C-D-E, while a sixth object was a sequence-irrelevant distractor. In each trial subjects chose objects by touching them and received either positive feedback (a yellow halo and high pitch sound) for correct, or negative feedback (a transient grey halo and low pitch sound) for choosing an object at an incorrect temporal position. After an erroneous choice, subjects had to re-choose the last correctly chosen object before making a choice of an object believed to be the next object in the sequence. Each trial allowed a maximum of ten errors to complete the sequence and receive fluid reward. If the sequence was not completed the objects were removed from the screen and a new trial was started with objects rearranged to new random locations. Subjects performed each sequence for a maximum of fifteen trials to allow for the gradual acquisition of the sequential ordering and additional trials to exploit the learned sequence. For each correctly chosen object, the position of the slider of a progress bar at the top of the screen stepped forward. Successful completion of a sequence always completed the slider progress bar and resulted in a fluid reward. When a sequence was not completed no fluid reward was given and the slider progress bar was reset for the subsequent trial with the same objects at new random locations. Choices of the distractor were registered as incorrect and not rewarded and required re-touching the last correctly chosen object.

### Hardware for automatic generation of Quaddle 2.0 objects

The Quaddle 2.0 toolbox has been developed and tested on an Apple MacOS system, but with minor adjustments it will be compatible with other operating systems that run Python and the Blender software (the tested Blender software version was 3.4.0). For the experiment, Quaddle 2.0 objects were generated with an Apple MacBook Pro 2023 with Apple M2 Max Chip and 32 GB Memory using MATLAB scripts (MATLAB 2022b) and python version 3.10.8. The objects presented here were generated with python scripts that connect with Blender version 3.4.0 (see **Figure 3**). Links to a comprehensive documentation and to the resources reproducing the objects are described in the Appendix.

## Results

### Automatized generation of multidimensional objects

The Quaddle 2.0 toolbox connects the object creation software Blender with MATLAB via a command-line interface, enabling users to generate objects bypassing Blender’s interface. This feature enables the automatized generation of large object databases with thousands of pre-configured objects. The MATLAB script *make_quaddle.m* (see **Appendix**) allows users to specify the number of objects, set the object similarity, and determine which features to keep or vary, providing high flexibility and utility for any application that requires multiple new objects across multiple cognitive experiments or augmented reality worlds. The script will generate an object table file for each specified object. A standard laptop is sufficient to generate objects. Objects displayed in Figure 1, 2, 4, and 6 were generated on an Apple MacBook Pro (2023) equipped with the Apple M2 Max chip and 32 GB of memory.

### Computing efficiency for different formats

The pre-configured scripts (see **Appendix**) allow users to select any combination of 3D object formats (FBX, GLTF) or a 2D image output (PNG). The time taken to generate each object varies depending on the selected output format. When generating GLTF files only, the average time to generate and save an object was 1.32 seconds per object. For FBX files only, the time increased slightly, averaging 1.55 seconds per object. Generating PNG images took longer, with an average time of 2.35 seconds per object. When generating GLTF, FBX, and PNG files simultaneously, the time increased to 2.42 seconds per object, primarily due to the additional time required to render the 2D image compared to saving 3D object formats directly.

### Improvements of the Quaddle 2.0 platform over previous versions

Quaddle 2.0 objects provide more feature dimensions and a larger feature space than Quaddle 1.0 objects (Watson et al., 2019), but can be used with less than ten feature dimensions, which allows reproducing objects with four feature dimensions similar to Quaddle 1.0 objects as shown in **Figure 1C**. The Quaddle 1.0 framework had other limitation that the new framework addresses. Quaddle 1.0 objects were rendered and generated from within the Studio DS Max software on windows PCs. By using the Blender software on PC or MacOS operating systems, and interfacing python scripts to Blender the Quaddle 2.0 platform moves beyond these limitations. This interface enables rapid ‘on-demand’ generation of objects for applications that require multiple new objects with random combinations of features as is typical for multiple types of learning studies that introduces novel objects over hundreds of experimental sessions (Banaie Boroujeni et al., 2022; Treuting et al., 2024; Womelsdorf, Watson, et al., 2021).

### Gradual morphing between features of multiple dimensions

Category learning studies have demonstrated that fine, gradual manipulations of object features is needed for controlling the discriminability of relevant dimensions in cognitive tasks (Apostel & Rose, 2021; Folstein et al., 2012; Minda & Smith, 2001). The Quaddle 2.0 platform enables such fine control of low level features and provides morphing functionality that can be applied to multiple features simultaneously as illustrated in **Figure 4**. Gradual changes of feature values include morphing *body shapes*, adjusting the *angle of stripe patterns*, and modifying *color shading*. Additionally, Quaddle 2.0 offers precise control over *arm angle* and the *bluntness or sharpness of arm tips* (**Figure 4**). Code and instructions for morphing specific color and pattern are available on the website (see **Appendix**).

### Quantitative evaluation of the similarity score of objects

The MATLAB object generation script *make_quaddle.m* (*see* Appendix) accompanying the Quaddle 2.0 toolbox calculates the similarity among objects by determining how many features are on average shared among objects in a given object set, weighting differently shared primary, secondary, and accessory features (*see* Methods). We quantitatively evaluated this similarity measure by comparing it with two existing alternative similarity measures. First, we calculated the Structural Similarity Index (SSIM), which measures the visual similarity between images by comparing luminance, contrast, and structure at the pixel level (Wang et al., 2004). We calculated the SSIM of Quaddle 2.0 objects exported as PNG images by applying the python package *skimage*. We computed the SSIM and the similarity score for 1000 pairs of object images randomly drawn from 300 randomly generated Quaddle 2.0 objects. Both, the SSIM and the similarity score were moderately positively correlated with similar average SSIM and similarity scores (SSIM: mean:0.4, variance: 0.03; Similarity Score: mean: 0.35, variance: 0.03) (**Figure 5A**). Secondly, we quantified how gradually increasing similarity scores of Quaddle 2.0 objects relate to a similarity score estimated by pretrained deep neural network models. These models process input data through multiple layers to extract meaningful patterns and features. At one or more hidden layers, the input data is transformed into dense vector representations known as feature embeddings. These embeddings are numerical outputs that capture the essential characteristics of images in a compact, multidimensional space (LeCun et al., 2015). We used the TensorFlow library (Abadi et al., 2016) to extract the feature vectors from the VGG16 model and the EfficientNetB0 model, which are pre-trained on large datasets like ImageNet (Simonyan & Zisserman, 2014; Tan & Le, 2019). Feature embedding vectors of two images were extracted and cosine similarity between two vectors were calculated as similarity score. The networks allow users to choose similarity metrics based on low-level structural comparisons or evaluation of high-level features-similarity. We found that the similarity score shows significant, albeit moderately positive correlations with the EfficientNetB0 similarity value with an *r*^2^ value of 0.09 (slope = 0.301; intercept = 0.332) (**Figure 5B**). Similarly, the VGG16 similarity value was also moderately correlated, with an *r*^2^ value of 0.127 (slope = 0.337; intercept = 0.255) (**Figure 5C**). These results show that scoring the similarity of Quaddle 2.0 objects based on shared feature spaces with a differential weighting of primary, secondary and minor feature dimensions is weakly but positively correlated with quantitative assessments of low-level feature similarity metrics.

### Empirical evaluation of Quaddle 2.0 objects and their similarity score

To test how subjects learn to associate different Quaddle 2.0 objects that are dissimilar and to test how higher similarity scores correspond to perceptual confusability of objects we tested rhesus monkeys on a sequence learning task. The testing was part of a larger empirical study (Wen et al., 2024) and included 125 testing sessions across four adult male rhesus monkeys (Subject B: 13, Subject J: 40, Subject K: 59, Subject S: 13). Each session contained on average 24 (Range: 9-26) sequence learning blocks. For each of the learning blocks on each of the sessions, six new Quaddle 2.0 objects were generated (five objects that were part of the sequence, and one distractor object). Subjects learned the predetermined 5-object sequences. They reached criterion performance (completing a sequence with maximally 15 choices, i.e. ten or less erroneous choices) within 6.53 ± 0.36 trials (subject B: 10.00 ± 1.39; J: 8.39 ± 0.58; K: 4.74 ± 0.40; S: 5.91 ± 0.89) (**Figure 6C**). Learning was accompanied by a steady decrease of the erroneous choices of the distractor (**Figure 6D**). In the experiment, the distractor object resembled either the second or fourth object of the sequence, and distractor errors occurred significantly more often at positions at which the sequence-relevant object and the distractor was similar, i.e. the distractor object was more easily confused with objects that had higher similarity scores (**Figure 6E**).

## Discussion

We introduced the Quaddle 2.0 toolbox for generating 3D rendered objects with multiple controllable feature dimensions for cognitive research and video gaming applications. Building on the foundation of the original Quaddle 1.0 framework, Quaddle 2.0 enhances stimulus generation by expanding number of independently varying feature dimensions and improving the script-based accessibility through the use of Blender, an open-source 3D modeling platform with Python support. Quaddle 2.0 allows using simple configuration text files to define objects and fine adjust low-level object features, which makes it easier to create custom object sets and automatize the generation of large object sets. Quaddle 2.0 supports various 3D model formats which enable their use in augmented and virtual reality platforms. Together, these toolbox features facilitate dynamic and immersive experimental designs.

It can replicate the key features from the original Quaddle framework (Watson et al., 2019), which has been successfully tested in experiments with both humans and non-human primates (Banaie Boroujeni et al., 2022; Boroujeni et al., 2022; Hassani et al., 2023; Kemp et al., 2024; Womelsdorf, Thomas, et al., 2021; Yazdanpanah et al., 2024). Extending this earlier framework, we have shown how the Quaddle 2.0 were used in sequence learning that required hundreds of new, unique objects in each experimental session, and which controlled the similarity among objects. Non-human primates learned to discriminate and serially order the objects and were confused by distractors when it shared a high feature similarity with target objects (**Figure 6**). Taken together, this example use case suggests that Quaddle 2.0 is a versatile, scalable, and user-friendly tool that enables cognitive research in 3D rendered environments.

## Appendix

- Python scripts and install instruction of the pipeline generating Quaddle 2.0 objects (shown in **Figure 3**) are available at https://github.com/xwen1765/blender-quaddle. Setting up the toolbox involves three steps: 1) Download Blender 3.4.0 and python code from the GitHub repository on the local computer. 2) Open Blender software, change preference as in the instruction (see below: *installation guide*). 3) Open the file *makeQuaddle.blend* and run the script parser.py, an example Quaddle will be created and can be viewed in Blender. To automatically generate object tables and generate 3D object sets with user defined similarity between objects, the user can use MATLAB. The GitHub repository includes the files *make_quaddle.m* and *test_example.m* (https://github.com/xwen1765/blender-quaddle/Matlab) that showcases how the configuration variables are set, object table files are generated and the parser.py is called to render and generate the Quaddle 2.0 objects.
- A website installation guide is available on https://xwen1765.github.io/posts/Quaddle/, video instruction is on https://www.youtube.com/watch?v=FOaKS-hQfYI.
- A website documentation about MATLAB scripts is available in the text file “*Documentation-MATLAB-make_quaddle.pdf*” on the website https://m-use.psy.vanderbilt.edu/quaddles-2/ and on the GitHub repository: https://github.com/xwen1765/blender-quaddle/Matlab/Documentation-MATLAB-make_quaddle.pdf
- An example of the morphing functionality is included in the *main.py* script on the GitHub repository. The script can be run directly to generate an example Quaddle in Blender. Users can specify parameters such as arm angle, body shape ratio, body pattern color, stripe angle, and beak size within the first few lines of the main function. Additionally, users can use the same function to add more arms (front and back) and other accessories beyond the default Quaddle feature set. To morph the pattern and color on the body, a Python script for generating pattern images is provided on the GitHub repository: https://github.com/xwen1765/blender-quaddle/PatternGeneration.

## Author Contributions

XW and TW were responsible for conceptualizing the research; XW developed the toolbox and designed the object features; XW and LM contributed programming code and generated the results and figures; TW supervised the study. All authors contributed to and have approved of the final version of the manuscript.

## Data availability

Not applicable.

## Code availability

All MATLAB and python scripts of the Quaddle 2.0 toolbox are publicly available via https://github.com/xwen1765/blender-quaddle

## Declarations

### Ethics approval

Not applicable.

### Consent to participate

Not applicable.

### Consent for publication

Not applicable.

